# Testing for heavy metals in drinking water collected from Dog Aging Project participants

**DOI:** 10.1101/2024.09.25.615013

**Authors:** Courtney L. Sexton, Janice O’Brien, Justin Lytle, Sam Rodgers, Amber Keyser, Mandy Kauffman, Matthew D. Dunbar, Dog Aging Project Consortium, Marc Edwards, Leigh Anne Krometis, Audrey Ruple

## Abstract

Heavy metals are commonly found in groundwater and can affect the quality of drinking water. In this pilot study, we analyzed the quality of drinking water for dogs participating in the Dog Aging Project (DAP) who lived in homes not served by a municipal water supply. In order to capture both diverse and localized environmental factors that may affect drinking water, 200 owners of DAP dogs located in one of 10 selected states were invited to participate. We tested for the presence of 28 metals in dogs’ drinking water, including eight (8) heavy metals that have maximum contaminant levels (MCLs) designated by the Environmental Protection Agency (EPA) and five (5) heavy metals that have EPA health guidance levels. The eight metals with MCLs are known to cause chronic health issues in humans after long-term ingestion. Our aim in this pilot was to determine whether such elements could be detected by at-home sampling of dogs’ drinking water. We found detectable levels of all metals tested. There were 126 instances when an analyte (arsenic, lead, copper, sodium, strontium, nickel, or vanadium) was above the EPA MCL or health guidance level. We further identified potential association between the presence of titanium and chromium, and occurrence of a known health condition in dogs. This prompts further investigation with a larger, stratified sample analyzing dogs’ drinking water composition and long-term health and wellness outcomes in dogs living in diverse geographies. These results may impact veterinary care decisions and husbandry, and underscore the validity and importance of utilizing dogs as sentinels of human health outcomes in the context of drinking water contamination.

## Introduction

Access to potable water is essential for animals’ survival, yet drinking water source, quality, and availability all vary greatly across species and populations [1–4]. For humans in the United States, drinking water has federally enforced minimum quality standards, per the Safe Drinking Water Act (SDWA) [5]. Federal water quality standards are monitored and implemented at the state level, and many states impose additional safety standards [6,7]. Despite substantial gains in access to safe drinking water since SDWA implementation [8,9], there is increasing concern that current infrastructure is vulnerable to changes in environmental factors, with high profile disasters such as the Flint, MI water crisis (2014), Gold King Mine disaster (2015) and train derailment in East Palestine, Ohio (2023) raising alarm regarding water and health [10–13]. Consumption of drinking water that contains elevated levels of metals, salts, bacteria, or other contaminants can lead to both acute and chronic health conditions, including cancer and organ failure in humans and dogs [14–20]. In addition to the contaminants currently regulated by the SDWA, the EPA maintains a Candidate Contaminant List (CCL) of additional contaminants, including several metals (Co, Li, Mn, Mo, V) that are of potential health concern, but whose toxicity profiles are not yet sufficiently definitive to derive standards [21].

Of concern, more than 15 million U.S. households rely on drinking water from privately owned wells, which are beyond the auspices of the SDWA. Because these systems operate on private land, maintenance and monitoring are solely the responsibility of the homeowner, and there are no reporting requirements. A growing body of literature has documented elevated levels of bacteria, lead, arsenic, radon, and other contaminants in drinking water collected from the point of use (POU) of homes dependent on private wells. As treatment of groundwater sourced from private wells prior to use is limited [22], there is concern that those dependent on these sources are uniquely vulnerable to exposure to heavy metals commonly found in groundwater [19] as well as to emerging anthropogenic contaminants [23]. People and animals who rely on unmonitored groundwater may be at higher risk of exposure to drinking water contaminated by sources including but not limited to landfill seepage, failed septic tanks, urban runoff, oil and gas extraction, and pesticide use [24–27]. Several recent studies have linked elevated rates of human illness to the consumption of contaminated well water [28–31].

For most companion animals, drinking water is provided by the people with whom they live and is typically from the same source (e.g. tap) those people use. Companion dogs typically have little awareness of the quality of or control over their drinking water sources and so may be especially susceptible to risks from chronic waterborne contaminants [32,33]. Previous and emerging research supports dogs’ role as sentinels of human health and wellbeing [34–36]; therefore, demonstration of adverse health effects related to waterborne consumption of heavy metals may identify critical health concerns for humans living in the same households as dogs [37] (Fig. 1).

**Figure 1.**
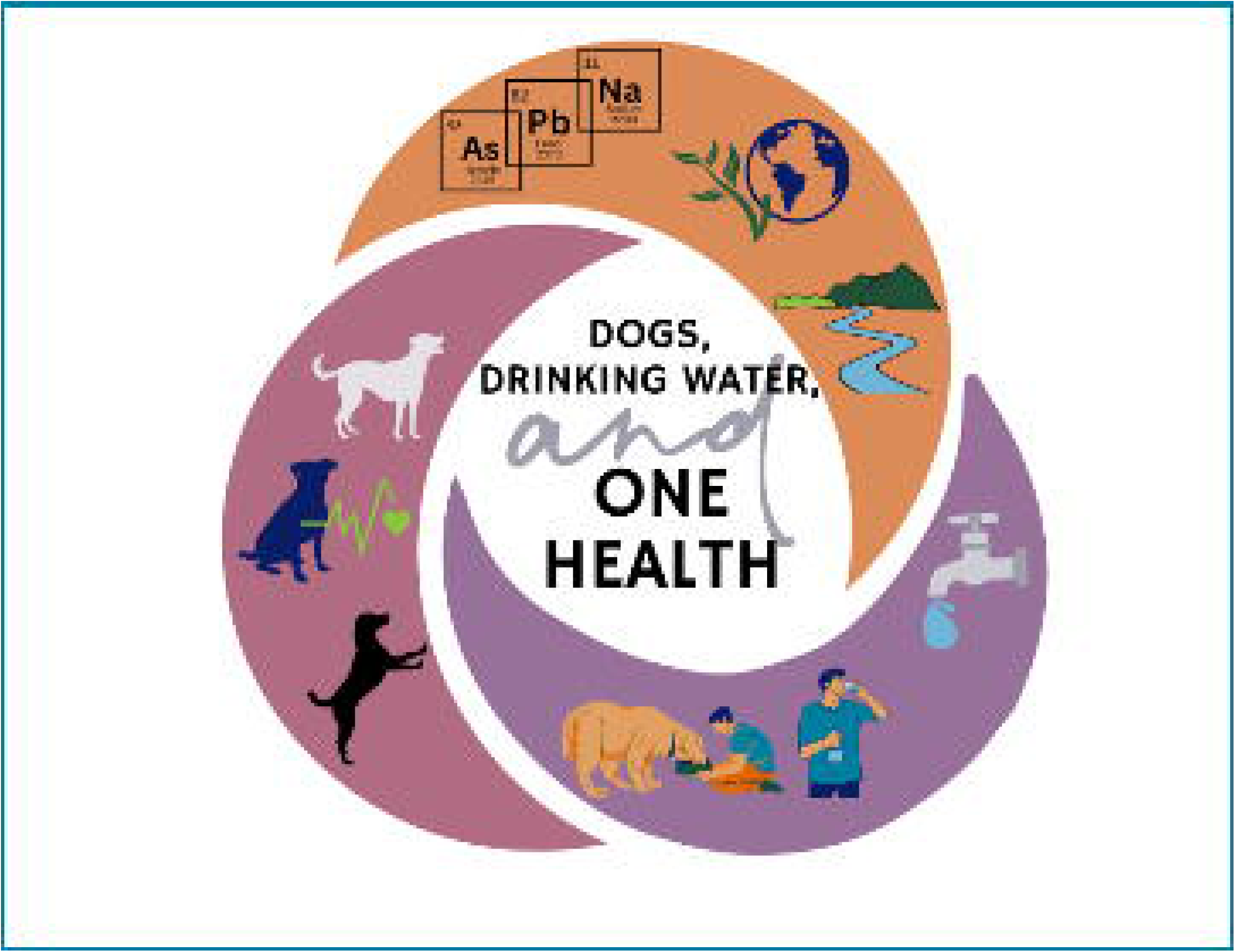
Companion dogs act as important sentinels of human health and wellbeing in considering a variety of risks and exposures, including potential illness from contaminated drinking water.

The aim of the present study was threefold: 1) to determine and quantify the presence of metals in drinking water samples intended for consumption by dogs; 2) to identify potential regional differences in the presence of various metals; and 3) to explore whether analyzing dogs’ drinking water could offer insight into potential sources of exposure-related outcomes, based on owner-reported information about their dogs’ health and wellbeing.

## Methods

### Recruitment & Sampling

The Dog Aging Project (DAP) is a longitudinal study of aging and age-related health outcomes in companion dogs [38]. Dog owners are recruited through various media outlets and word of mouth to voluntarily nominate their dogs for participation. Owners who opt to enroll their dogs in the study are first led through an extensive informed consent process. Once consent is confirmed, owners are requested to complete the Health and Life Experiences Survey (HLES), which includes a number of small surveys that collect information about dog demographic characteristics, physical activity, environment, dog behavior, diet, medications and preventatives, dog health status and history, and owner demographic characteristics. Completion of HLES defines enrollment in the DAP Pack. There are additional informed consent processes for dogs selected to participate in additional sampled cohorts, which may involve collecting such data as electronic veterinary medical records, genome-wide sequence information, clinicopathology and molecular phenotypes derived from blood cells, plasma, and fecal samples. The project began enrollment in 2019 and had over 47,000 dogs enrolled as of January 1, 2023.

For the present study, we recruited a subset of DAP participants who previously reported having a non-municipal water supply. In order to capture both diverse and localized environmental factors that may affect drinking water, 200 owners of DAP dogs located in one of 10 selected states (California, Florida, Michigan, North Carolina, New York, Pennsylvania, Texas, Virginia, Washington, and Wisconsin) were invited to participate.

Dog owners completed a pre-sampling survey (see supplementary materials) in which they described the source and quality of their dog’s drinking water. Once they completed the survey, owners were sent kits via mail that included sealed water bottles and instructions for collecting water samples from their dog’s primary drinking water source (see supplementary materials).

Participants collected samples from their dog’s primary drinking water source at a single time point after a minimum of 6 hours stagnation (lack of water use) in labeled collection vials, and returned the samples to our lab via prepaid mailers.

### Ethics Statement

The University of Washington IRB deemed that recruitment of dog owners for the Dog Aging Project is human subjects research that qualifies for Category 2 exempt status (IRB ID no. 5988, effective 10/30/2018). All study-related procedures and sample collection involving privately owned dogs were approved by the Texas A&M University IACUC, under AUP 2021-0316 CAM (effective 12/14/2021).

### Sample Analyses

All water samples were acidified to 2% nitric acid (v/v) and digested for 16 hours before analysis for metals concentrations by a Thermo Electron X-Series ICP-MS per Standard Method 3125 B [39]. Blanks and spikes of known metal concentrations were processed for each sampling event for QA/QC purposes. The presence and quantity of 28 different metals and minerals were determined for each water sample, including eight heavy metals – chromium, copper, arsenic, selenium, cadmium, barium, lead, and uranium – with EPA-designated maximum contamination level (MCL) goals; five heavy metals – sodium, strontium, nickel, vanadium, and cobalt – with EPA-designated health guidance levels; and six elements – aluminum, copper, iron, manganese, silver, and zinc – with secondary maximum contaminant levels (SMCLs) relevant to taste, color, and odor. The minimum reporting level (MRL) for detection of each of the 28 metals and minerals can be found in Table S1.

### Data Return

Once analyses were completed, we reported back to dog owners via email the raw data regarding the quantities of all 28 heavy metals and minerals detected in their dog’s drinking water sample, as well as links to additional information from the EPA regarding the impacts of heavy metal exposure on health and how to safely control levels of these metals. Data returns included a graphic scale illustrating how levels of each of the eight metals with EPA designated health action levels (MCLs) in their dog’s drinking water compared to the upper limit. For the additional 20 metals without MCLs tested, data returns included a table listing each element with results in either parts per million (ppm) or parts per billion (ppb). In the absence of known MCLs for these metals, we provided the quartile in which participants’ results fell compared to other study participants (see supplementary materials). If a participant dog’s drinking water sample showed a heavy metal level of more than four times the EPA MCL, the participant was immediately notified of this incidental finding via phone call or voice message, with a follow-up email providing additional information including links to EPA websites to provide the dog’s owner with more information about how to mitigate further exposure.

### Statistical Analyses

The minimum reporting level (MRL) is the minimum concentration that can be reported by a laboratory as a quantified value for a method analyte in a sample following analysis. Due to the nature of the testing method, some individual results fall below the MRL. In our statistical analyses, these were coded as 0 as they were reasonably close to 0. For each of the 28 individual metals or minerals, the medians, means, standard deviations, and ranges were computed using these quantified values.

For each metal with an EPA-designated MCL, SMCL, and guidance level [5], the proportion of samples that exceeded the relevant EPA level were calculated for each individual metal. Additionally, the total number of samples that exceeded any MCL or guidance level (but not sMCL) were calculated. For this calculation, if a sample exceeded the limit in any metal, or even in multiple metals, it was considered over the limit. Samples that exceeded multiple metal limits were not counted twice.

Regarding water treatment systems, each system type was categorized according to intended purpose, including a) health intervention (ultraviolet, reverse osmosis, chlorination, carbon filter); or b) aesthetic intervention (acid neutralizer, sediment filter, water softener, iron removal). For each respondent, the number of unique installed systems was tabulated— e.g., if a respondent had a sediment filter and an ultraviolet light system installed, they would have two systems installed. Participants were then grouped according to whether they reported using systems categorized as health intervention only, both health and aesthetic intervention, aesthetic intervention only, other, none (for no installed system) and not sure (respondents unsure if an installed system was present).

#### Regression analyses

Several regression models were built in R studio version 4.4.0 using the glm function in order to examine associations. The first group of models investigated associations between multiple water source variables as reported by owners in the pre-sampling survey and the quantification of each metal. Responses that included “unknown” or “other” for any of the variables were excluded from the analysis. The input variables were coded as either binary, or categorical/ordinal (depending on the number of survey response options). The frequency of pest and weed treatment were treated as ordinal variables. The input variable for well age was binary with options set as before and after the population median well install year of 1996. The glm family used in the analysis was Gaussian as the analyte output values were numeric. The water source variables that correlated with statistical significance (p<0.1) were recorded for each metal. A more lenient alpha cutoff was used in this analysis as this was a hypothesis-generating examination where identifying preliminary evidence of associations was of higher importance.

A second set of analyses investigated possible associations between numerical metal value and the total burden of developed disease for each dog. The total number of owner-reported diseases that were non-congenital were calculated for each dog. The glm family used in the analysis was Poisson, as the disease numbers were counts. The input was the numerical value for all the metals, and the output was the disease count. Age was added to the model to control for potential confounding, as age is associated with disease burden. The coefficients were exponentiated to calculate the rate ratio.

The third model assessed possible associations between the water treatment system installed and the total burden of developed disease. The glm family used in the analysis was Poisson. The input was the binary value for each treatment system (whether that system was installed yes/no) with “none” as its own category of system, and the output was the developed disease count. Age was added to the model to control for potential confounding. The coefficients were exponentiated to calculate the rate ratio.

## Results

A total of 178 of the 200 (89.0%) sample kits sent were returned by dog owners. In seven cases, kits were lost in the mail either on the way to participants or after they had been sent back to our lab for analysis. In two cases, dogs had passed away before sampling occurred and the owners opted out of testing after the sampling kit had been deployed.

### Dog-Owner Demographics

There were at least 16 participants from each of the 10 selected states. The majority (97%) of dog owners identified as White. Owners ranged in age group from 18-24 to >75, with 63% of owners being aged 55 or older. Fifteen percent of owners had a reported income of < $60K, and 74% had received at least some amount of higher education. Participants’ home locations were predominantly located in rural (72%) locations, with a minority (28%) located in suburban or urban areas; this is to be expected, given that private wells are most common in rural areas [40]. Most (87%) owners reported their dog to be in very good to excellent health, even though owners of 147/178 dogs (82%) reported their dog as having a known developed (i.e., non-congenital) health condition of some kind (Table 1).

**Table 1.** Dog-owner pair demographic information for all participants who returned a water sample kit.

|  |  | COUNT (N= 178) | PERCENT |
| --- | --- | --- | --- |
| OWNER IDENTIFIED RACE | Alaska Native | 0 | 0.0 |
|  | American Indian | 3 | 1.7 |
|  | Asian | 3 | 1.7 |
|  | Black or African American | 0 | 0.0 |
|  | Native Hawaiian | 0 | 0.0 |
|  | Other | 4 | 2.2 |
|  | Other Pacific Islander | 0 | 0.0 |
|  | White | 173 | 97.0 |
| OWNER AGE GROUP | 18-24 | 2 | 1.1 |
|  | 25-34 | 14 | 7.9 |
|  | 35-44 | 18 | 10.1 |
|  | 45-54 | 32 | 18.0 |
|  | 55-64 | 59 | 33.0 |
|  | 65-74 | 40 | 22.5 |
|  | >75 | 14 | 7.9 |
| OWNER INCOME LEVEL | <20k | 3 | 1.7 |
|  | 20-39k | 12 | 6.7 |
|  | 40-59k | 12 | 6.7 |
|  | 60-79k | 18 | 10.1 |
|  | 80-99k | 24 | 13.5 |
|  | 100-119k | 20 | 11.2 |
|  | 120-139k | 13 | 7.3 |
|  | 140-159k | 7 | 3.9 |
|  | 160-179k | 6 | 3.4 |
|  | >180k | 36 | 20.2 |
|  | not say | 28 | 15.7 |
| **OWNER HIGHEST EDUCATION LEVEL** | HS Grad | 5 | 2.8 |
|  | GED or Equivalent | 3 | 1.7 |
|  | Some College, No Degree | 20 | 11.2 |
|  | Trade or Vocational | 7 | 3.9 |
|  | Associates | 12 | 6.7 |
|  | Bachelors | 50 | 28.0 |
|  | Masters | 57 | 32.0 |
|  | Professional | 13 | 7.3 |
|  | Doctorate | 12 | 6.7 |
| **STATE** | CA | 17 | 9.6 |
|  | FL | 19 | 10.7 |
|  | MI | 17 | 9.6 |
|  | NC | 17 | 9.6 |
|  | NY | 16 | 9.0 |
|  | PA | 18 | 10.1 |
|  | TX | 18 | 10.1 |
|  | VA | 20 | 11.2 |
|  | WA | 17 | 9.6 |
|  | WI | 20 | 11.2 |
| **HOME LOCATION CLASSIFICATION** | Rural | 129 | 72.5 |
|  | Suburban | 45 | 25.3 |
|  | Urban | 5 | 2.8 |
| **OWNER REPORTED DOG HEALTH STATUS** | Excellent | 100 | 56.0 |
|  | Very Good | 56 | 31.4 |
|  | Good | 15 | 8.4 |
|  | Fair | 6 | 3.4 |
|  | Poor | 1 | 0.6 |
|  | Very Poor | 1 | 0.6 |
| **OWNER REPORTED DEVELOPED HEALTH CONDITIONS IN THE DOG PARTICIPANT (I.E. NON-CONGENITAL)** | No developed conditions | 32 | 17.9 |
|  | Developed conditions | 147 | 82.6 |
| **DOG AGE AT TIME OF STUDY** | 0-1 yr | 11 | 6.2 |
|  | 2-4 yr | 36 | 20.2 |
|  | 5-9 yr | 83 | 46.6 |
|  | 10+ yr | 50 | 28.0 |
| **DOG BREED STATUS** | Purebred | 99 | 55.6 |
|  | Mixed breed | 81 | 45.5 |
| **DOG WEIGHT (KG)** | 0-9.9 | 34 | 19.1 |
|  | 10-19.9 | 30 | 16.9 |
|  | 20-29.9 | 60 | 33.7 |
|  | 30-39.9 | 32 | 18.0 |
|  | 40+ | 24 | 13.5 |

### Household Water Survey

All dog owners who returned a kit responded to a pre-sampling survey in which they described their understanding of their dogs’ water supply, reporting water source, year the well was constructed, potential leaching sources within 100ft of the water supply, geographic features within 0.5 miles of the water source, types of installed treatment systems (if any), water testing history, composition of water pipes within the home, water density, weed or pest treatment, and any discernible abnormalities in taste, smell or color; survey questions were adapted from a successful instrument employed by Cooperative Extension and collaborators for over ten years [41]. These results are summarized in Table 2, with additional survey information available in the supplementary materials.

**Table 2.** Water Source Variables. Detailed information about participant dogs’ drinking water sources as reported by dog owners (N=178).

|  |  | COUNT (N=178) | PERCENT |
| --- | --- | --- | --- |
| WATER SOURCE | Treated | 117 | 65.7 |
|  | Untreated | 57 | 32 |
|  | Unknown/Other | 4 | 2.2 |
| **WELL YEAR** | Before 1996 | 58 | 32.6 |
|  | 1996 → | 40 | 22.5 |
|  | Unknown | 80 | 44.9 |
| **REPORTED POTENTIAL LEACHING SOURCES (WITHIN 100 FT) <sup>x</sup>** | None | 140 | 78.7 |
|  | Potential Source of Leaching <sup>x</sup> | 29 | 16.2 |
|  | Unknown | 9 | 5.1 |
| **REPORTED GEOGRAPHIC FEATURES WITHIN 0.5 MILE <sup>∞</sup>** | None | 104 | 58.4 |
|  | Geographic Feature <sup>∞</sup> | 63 | 35.4 |
|  | Unknown | 11 | 6.2 |
| **INSTALLED HOME WATER SYSTEMS <sup>ε</sup>** | None | 50 | 28 |
|  | Installed System - Aesthetic <sup>ε</sup> | 66 | 37 |
|  | Installed System - Health <sup>ε</sup> | 23 | 12.9 |
|  | Installed System - Both <sup>ε</sup> | 32 | 18 |
|  | Unknown | 8 | 45 |
| **EVER HAD WATER TESTED** | Yes | 116 | 65.1 |
|  | No | 62 | 34.8 |
| **PIPES TYPE** | Plastic | 117 | 65.7 |
|  | Metal | 23 | 13 |
|  | Unknown | 27 | 15.2 |
|  | Did Not Report | 11 | 6.2 |
| **PIPES ISSUES** | Yes | 9 | 5.1 |
|  | No | 169 | 94.9 |
| **HARD OR SOFT WATER (N=176)** | Hard | 89 | 50.5 |
|  | Soft | 40 | 22.7 |
|  | Neither | 36 | 20.5 |
|  | Unknown | 11 | 6.3 |
| **WEED TREATMENT** | Regular | 57 | 32 |
|  | Sporadic | 25 | 14 |
|  | Never | 96 | 53.9 |
| **PEST TREATMENT** | Regular | 30 | 16.9 |
|  | Sporadic | 39 | 21.9 |
|  | Never | 109 | 61.2 |
| **APPARENT ABNORMALITIES <sup>°</sup>** | Yes <sup>°</sup> | 54 | 30.3 |
|  | No | 124 | 69.7 |

<sup>x</sup> Leaching sources include: septic drain field, pit privy or outhouse, home heating oil storage tank, pond or freshwater stream, cemetery, tidal shoreline/estuary/marsh.
<sup>∞</sup> Geographic features include: Landfill, golf course, abandoned quarry or industrial site, active quarry, active industrial/manufacturing/processing site, dump site, field crops/plant nursery, farm animal operation, railroad tracks, airport, fracking site, commercial underground storage tank or supply lines.
<sup>ε</sup> Installed home water systems include: Health-based – UV light, activated carbon/charcoal filter, reverse osmosis, chlorination system; Aesthetic – sediment filter, acid neutralizer, water softener/conditioner, iron removal.
<sup>°</sup> Apparent abnormalities include staining on fixtures, unpleasant taste, unpleasant odor, unnatural appearance, and particles visible.

One-third of owners were served by wells > 25 years old (33%), and 117 owners (65%) reported having their well water treated. A majority of people (68%) further installed some form of water treatment system within their home. Of these, 13% installed health-based systems including UV light, activated carbon/charcoal filter, reverse osmosis, and chlorination systems. Another 37% installed aesthetic-based systems (e.g., to alter smell/taste/appearance) including sediment filters, acid neutralizer, water softener/conditioner, and iron removal. An additional 18% of participants reported having installed both aesthetic- and health-based systems (Fig. 2).

**Figure 2.**
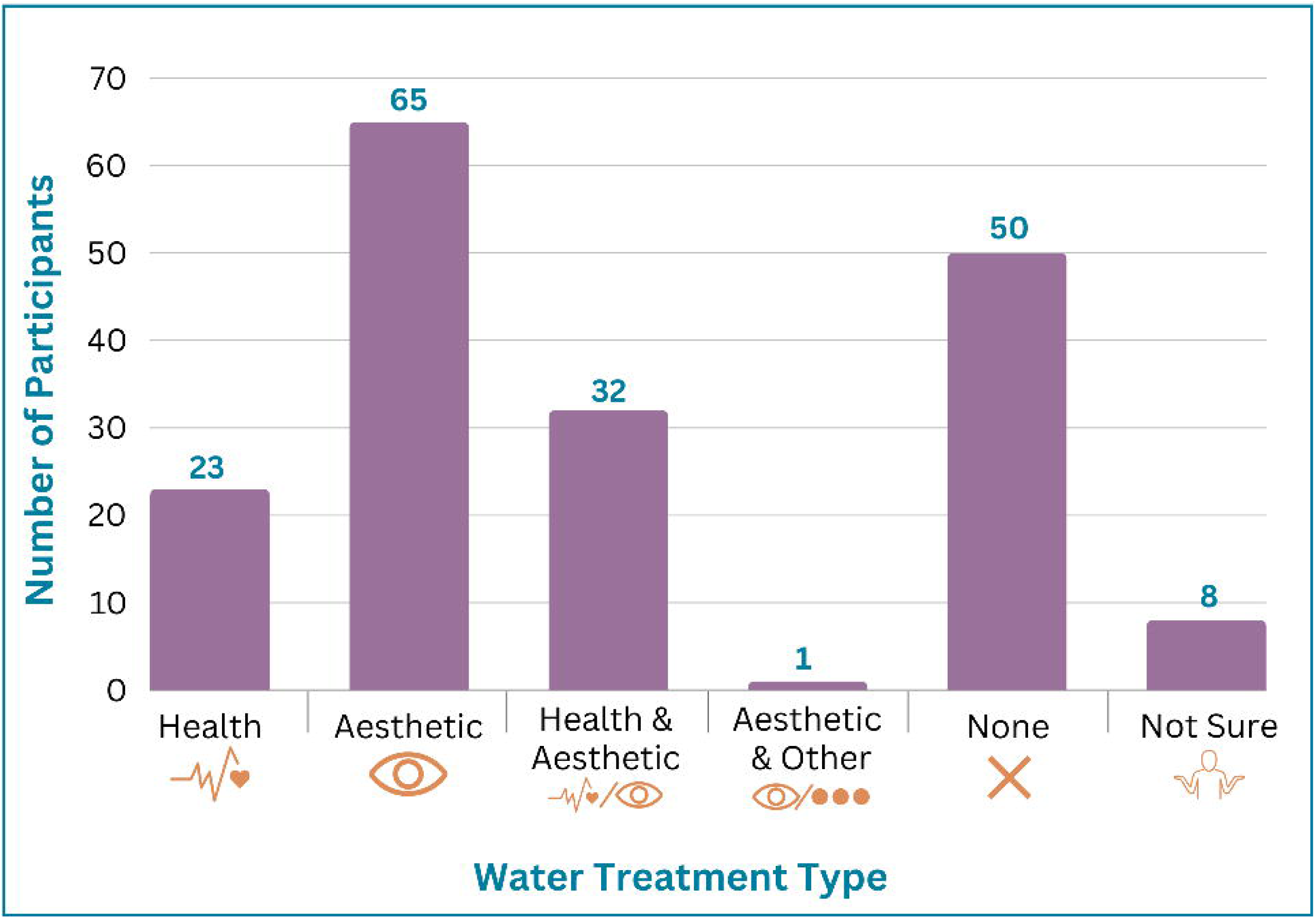
Dog owners reported using a variety of home filtration systems, with just over half (55%) installing health-based systems.

Twenty-nine owners (16%) reported an awareness of a potential leaching source (e.g., septic drain field, pit privy or outhouse, home heating oil storage tank, pond or freshwater stream, cemetery, or tidal shoreline/estuary/marsh) being located within 100 feet of their well. One hundred four (58%) identified certain geographic features within 0.5 miles of their well (e.g., landfill, golf course, abandoned quarry or industrial site, active quarry, active industrial/manufacturing/processing site, dump site, field crops/plant nursery, farm animal operation, railroad tracks, airport, fracking site, commercial underground storage tank or supply lines). Eighty-seven people (49%) reported regular use of pest or weed treatment on their property.

One hundred seventeen people (66%) reported having plastic water pipes, and another 23 (13%) reported metal pipes within their home. Twenty-seven (15%) did not know their pipe material. Half of participants (89) reported having “hard” water, compared to 22% (40) who reported “soft” water, and 21% (36) did not know. Nine people (5%) reported having pipe issues. Approximately one-third (54) reported apparent abnormalities related to their well water, such as staining on fixtures, unpleasant taste, unpleasant odor, unnatural appearance, and visible particles.

### Water Quality Analyses

Regarding overall detection levels, total quantities of analytes detected varied substantially from sample to sample, with respect to each metal and mineral (Table 3; Table S1), and according to geographic location (Fig. 3). All analytes were above the MRL in at least one instance. Twenty-one elements (75% of those analyzed), including chromium, copper, barium, lead, uranium, lithium, aluminum, calcium, chlorine, cobalt, iron, manganese, nickel, phosphorus, potassium, strontium, sulfur, tin, titanium, vanadium, and zinc were above the MRL in at least one sample in every state included in this pilot.

**Figure 3.**
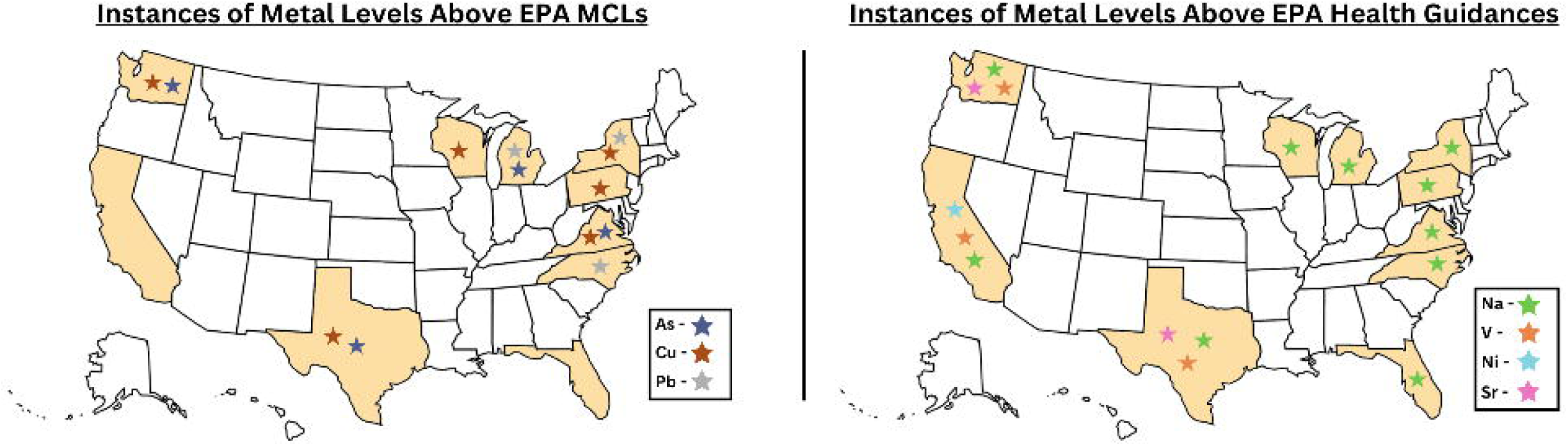
Copper and/or lead and/or arsenic levels were above the EPA maximum contaminant level (MCL) in eight out of ten states. Sodium, strontium, nickel and vanadium surpassed the health guidance level in several states.

**Table 3.** Sample variance of metals tested in dog drinking water samples, per EPA action and guidance levels (ppm).

| MEASURE OF WATER QUALITY | STANDARD OR RECOMMENDATION | % EXCEEDANCE | ELEMENT MEDIAN (MEAN ± DEVIATION) | ELEMENT MAX |
| --- | --- | --- | --- | --- |
| **EPA MCLs/ACTION LEVEL (HEALTH)** |  |  |  |  |
| **COPPER** | <1.3 ppm | 3.4% | 0.045 (0.19±0.35) | 2.4 |
| **ARSENIC** | <0.010 ppm | 2.2% | 0.0002 (0.0013±0.0053) | 0.066 |
| **LEAD** | <0.015 ppm | 1.7% | 0.0005 (0.0016±0.0033) | 0.022 |
| **BARIUM** | <2 ppm | 0% | 0.0054 (0.024±0.043) | 0.28 |
| **CADMIUM** | <0.005 ppm | 0% | 0 (0.000035±0.00021) | 0.0027 |
| **CHROMIUM** | <0.1 ppm | 0% | 0.0007 (0.001±0.0013) | 0.013 |
| **SELENIUM** | <0.050 ppm | 0% | 0.0003 (0.00088±0.0032) | 0.03 |
| **URANIUM** | <0.030 ppm | 0% | 0.0002 (0.0010±0.0035) | 0.03 |
| **EPA SMCLs/ACTION LEVEL (TASTE AND AESTHETICS)** |  |  |  |  |
| **ALUMINUM** | <0.05 ppm | 22% | 0.0064 (0.067±0.22) | 2.3 |
| **MANGANESE** | <0.05 ppm | 7.8% | 0.0009 (0.023±0.099) | 0.85 |
| **COPPER** | <1 ppm | 3.9% | 0.045 (0.19±0.35) | 2.4 |
| **IRON** | <0.3 ppm | 1.7% | 0.0078 (0.040±0.15) | 1.9 |
| **SILVER** | <0.1 ppm | 0% | 0 (0.0000050±0.000035) | 0.0003 |
| **ZINC** | <5 ppm | 0% | 0.064 (0.26±0.55) | 4.3 |
| **EPA GUIDANCE/HEALTH REFERENCE LEVELS** |  |  |  |  |
| **SODIUM** | <20 ppm | 59% | 31 (63±77) | 450 |
| **VANADIUM** | <0.021 ppm | 2.2% | 0.0003 (0.0023±0.0082) | 0.074 |
| **STRONTIUM** | <4 ppm | 1.1% | 0.052 (0.26±0.76) | 7.9 |
| **NICKEL** | <0.1 ppm | 0.6% | 0.001 (0.0062±0.047) | 0.63 |
| **COBALT** | <0.07 ppm | 0% | 0.0001 (0.00014±0.00032) | 0.0024 |

Regarding health and guidance levels, in total, there were 126 instances of an analyte being above an EPA MCL or guidance level— with 114 dogs’ (64%) samples with at least one metal being above an EPA MCL or guidance level. There were 13 instances (7% of samples) where values of arsenic, lead, or copper were above the EPA MCLs. These samples were located in 8/10 states included in this study (Fig. 3). Additionally, there were 106 instances (60% of samples) in which sodium was above the EPA health guidance level, ranging across every state sampled. There were several instances among our samples where other metals with health guidance levels also surpassed these guidelines, broken down as follows: vanadium – 4 instances (in CA, TX, and WA); strontium – 2 instances (in TX and WA); nickel – 1 instance (in CA) (Fig. 3).

### Regressions

#### Qualitative Characteristics

We found several statistical associations between features of dogs’ water sources and various metals (Table 4). On average, wells installed after 1996 had lower values of silicon [-1277, SE = 721, p=0.08], copper [-79, SE = 36, p=0.03], zinc [-140, SE = 71, p=0.05], and lead [-0.71, SE = 0.3, p=0.02] compared to older wells.

**Table 4:**
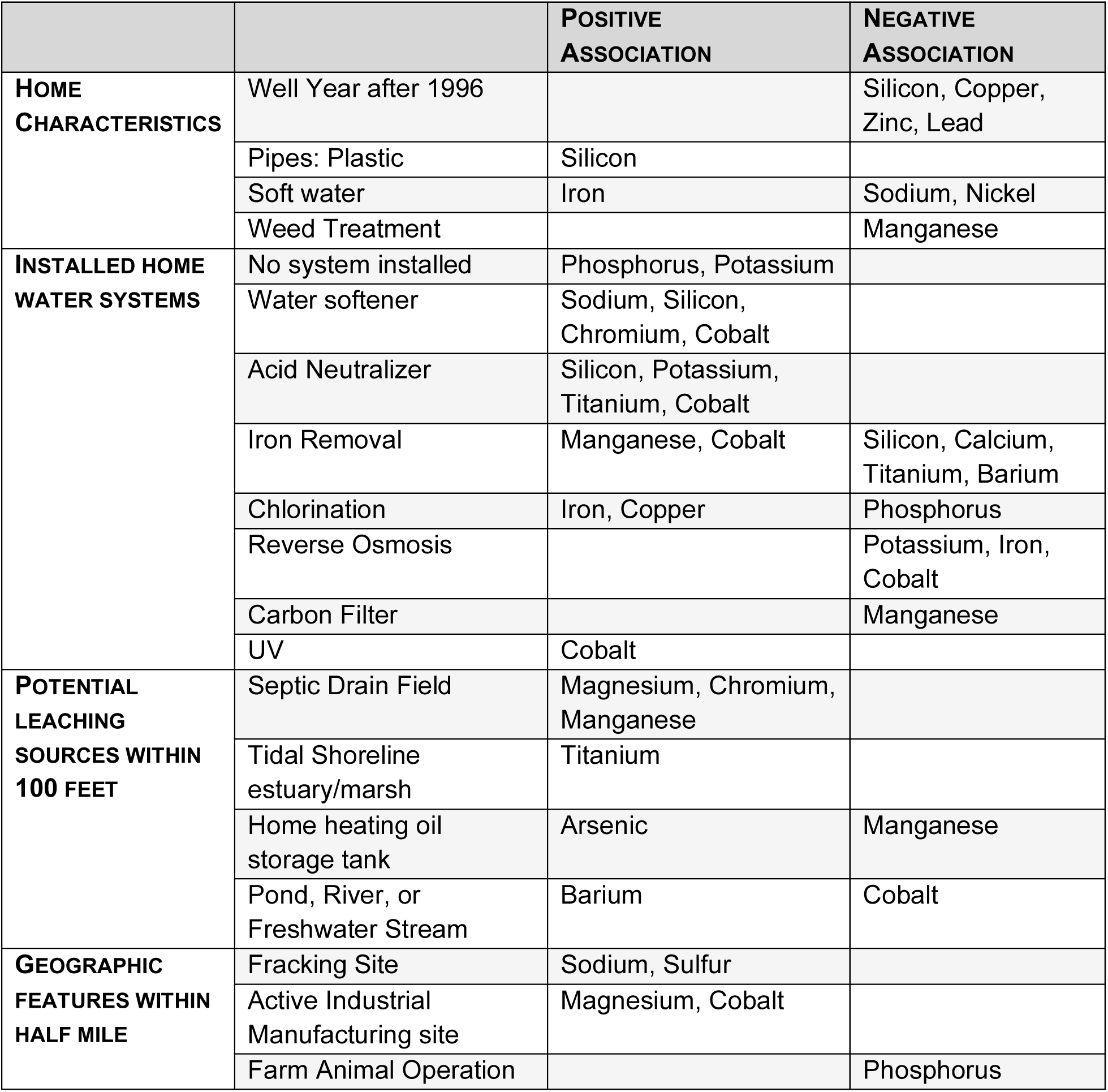
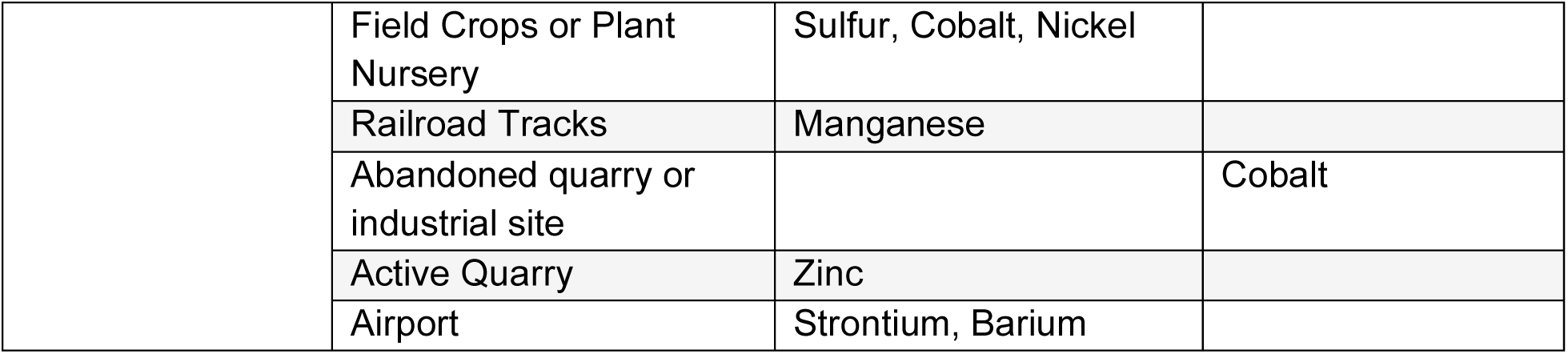
Water source variables and statistically significant (p<0.1) associations with individual metal levels.

Wells located within 100 feet of potential leaching sources were also associated with different metals depending on the potential leaching source. Septic drain fields positively correlated with magnesium [12717, SE = 6157, p=0.04], chromium [1.1, SE = 0.6, p=0.05], and manganese [214, SE= 51, p=9e-05]. The presence of tidal shoreline/estuary/marsh positively associated with titanium values [1.8, SE = 1.0, p=0.08]. Home heating oil storage tanks positively associated with arsenic values [5.4, SE = 1.2, p=4e-05], but negatively with manganese [-258, SE = 73, p=0.0008]. Being next to a pond, river, or freshwater stream positively correlated with barium [30.7, SE = 18, p=0.093], but negatively associated with cobalt [-0.3, SE = 0.1, p=0.06].

Wells within a 0.5 mile of various geographic features associated with different metals depending on the feature. Reported fracking sites positively associated with sodium [175167, SE = 61721, p=0.006] and sulfur [86, SE = 46, p=0.06]. Active industrial manufacturing sites positively associated with magnesium [37099, SE = 14796, p=0.01] and cobalt [1.32, p=0.0009]. Farm animal operations negatively associated with phosphorus results [-51.5, p=0.08]. Field crops and plant nurseries positively associated with sulfur [24.9, p=0.07], cobalt [0.15, SE = 0.4, p=0.06], and nickel [36, SE = 17, p=0.04] concentrations. Railroad tracks positively associated with manganese values [370.4, SE = 79, p=1.52e-05]. Abandoned quarries negatively associated with cobalt values [-1.2, SE = 0.6, p=0.03], while active quarries positively associated with zinc [1099.5, SE = 568, p=0.06]. Airports positively correlated with the levels of strontium [1070.3, SE = 401, p=0.009] and barium [142.7, SE = 43, p=0.001].

Water reported as “soft” water positively correlated with levels of iron in water [28.2, SE = 16, p=0.08], and negatively with nickel [-45.8, SE = 22, p=0.04] and sodium [-48881.1, SE = 23599, p=0.04]. Plastic pipe material was positively associated with silicone [4417.2, SE = 2606, p=0.09]. Metal pipes did not associate with any metal presence in the water samples.

Both regular and sporadic use of weed treatment correlated negatively with manganese [sporadic -77.8, SE = 33, p=0.02; regular -115.4, SE = 43, p=0.009].

#### Health Outcomes

Regarding the presence of individual metals, titanium (RR 0.81, 0.68-0.97) and chromium (RR 0.87, 0.76-0.97) were associated with developed health outcomes (Table 5). They were both below a rate ratio of 1.0, meaning that higher values of these elements correlated with fewer health diagnoses. The remainder of the metals did not statistically significantly associate with reported health outcomes in our sample.

**Table 5:** Computed rate ratio estimates for each metal level and owner-reported non-congenital (developed) health outcomes in dogs.

|  | **RATE**<br>**RATIO** | **95% CI**<br>**LOW** | **95% CI**<br>**HIGH** | **P-**<br>**VALUE** |
| --- | --- | --- | --- | --- |
| **LITHIUM** | 0.99 | 0.98 | 1.00 | 0.06 |
| **SODIUM** | 1.00 | 1.00 | 1.00 | 0.18 |
| **MAGNESIUM** | 1.00 | 1.00 | 1.00 | 0.20 |
| **ALUMINUM** | 1.00 | 1.00 | 1.00 | 0.02 |
| **SILICON** | 1.00 | 1.00 | 1.00 | 0.17 |
| **PHOSPHORUS** | 1.00 | 1.00 | 1.00 | 0.35 |
| **SULFUR** | 1.00 | 0.99 | 1.00 | 0.63 |
| **CHLORINE** | 1.00 | 1.00 | 1.00 | 0.29 |
| **POTASSIUM** | 1.00 | 1.00 | 1.00 | 0.06 |
| **CALCIUM** | 1.00 | 1.00 | 1.00 | 0.52 |
| **TITANIUM** | 0.81 | 0.68 | 0.97 | 0.02* |
| **VANADIUM** | 0.99 | 0.97 | 1.01 | 0.49 |
| **CHROMIUM** | 0.87 | 0.76 | 0.97 | 0.02* |
| **IRON** | 1.00 | 1.00 | 1.00 | 0.36 |
| **MANGANESE** | 1.00 | 1.00 | 1.00 | 0.50 |
| **COBALT** | 0.80 | 0.55 | 1.09 | 0.19 |
| **NICKEL** | 1.00 | 0.99 | 1.00 | 0.37 |
| **COPPER** | 1.00 | 1.00 | 1.00 | 0.48 |
| **ZINC** | 1.00 | 1.00 | 1.00 | 0.16 |
| **ARSENIC** | 1.00 | 0.98 | 1.02 | 0.73 |
| **SELENIUM** | 0.98 | 0.93 | 1.02 | 0.36 |
| **STRONTIUM** | 1.00 | 1.00 | 1.00 | 0.98 |
| **SILVER** | 0.82 | 0.05 | 10.81 | 0.89 |
| **CADMIUM** | 1.21 | 0.41 | 3.95 | 0.75 |
| **TIN** | 2.03 | 0.57 | 6.48 | 0.25 |
| **BARIUM** | 1.00 | 1.00 | 1.00 | 0.31 |
| **LEAD** | 0.96 | 0.92 | 1.00 | 0.08 |
| **URANIUM** | 1.00 | 0.96 | 1.05 | 0.91 |

For reported presence of home water treatment systems, which are considered health interventions, we found that the presence of a reverse osmosis system was negatively associated with the number of diagnosed health conditions (RR = 0.7). In other words, the statistical association suggests dogs drinking water that was treated by reverse osmosis were less likely to have a diagnosed health condition (Table 6).

**Table 6:** Computed rate ratio estimates for different types of installed water systems and owner-reported non-congenital (developed) health outcomes in dogs.

|  | **RATE**<br>**RATIO** | **95% CI**<br>**LOW** | **95% CI**<br>**HIGH** | **P-**<br>**VALUE** |
| --- | --- | --- | --- | --- |
| **NONE** | 1.09 | 0.84 | 1.41 | 0.52 |
| **UV** | 1.20 | 0.86 | 1.65 | 0.27 |
| **WATER SOFTENER** | 1.23 | 0.99 | 1.53 | 0.06 |
| **SEDIMENT FILTER** | 1.39 | 1.14 | 1.70 | <0.001* |
| **REVERSE OSMOSIS** | 0.70 | 0.51 | 0.93 | 0.02* |
| **IRON REMOVAL** | 0.84 | 0.62 | 1.12 | 0.24 |
| **ACTIVATED CHARCOAL FILTER** | 1.12 | 0.88 | 1.42 | 0.34 |
| **CHLORINATION** | 0.97 | 0.64 | 1.42 | 0.88 |
| **ACID NEUTRALIZER** | 1.04 | 0.58 | 1.75 | 0.88 |

For reported presence of water treatment systems considered aesthetic, we saw the following associations with reported health outcomes in dogs: Sediment filters positively correlated with the number of diagnosed health conditions (RR = 1.39, 1.14-1.7). In other words, dogs drinking water that was treated with a sediment filter were more likely to have diagnosed health conditions. Water softeners positively correlated as well (RR = 1.23, 0.99-1.53), however even though there was weak evidence (p = 0.06), the rate ratio estimate crossed 1 (Table 6).

## Discussion

Our results confirm that a variety of heavy metals are detectable in well water via direct sampling of dogs’ drinking water. Local land use and household factors influence the levels of metals detected, and there is geographic variability in the types/levels of metals detected.

The incidence of metals contamination and maximum observed levels are noticeably lower than in previous studies examining point of use water quality in private wells [30,42–46]. This may be because the average household income and education level was high, which is often associated with better in-home water quality [22]. Regardless, even in this ‘best case scenario’, three heavy metals (arsenic, lead, and copper) were detected in samples submitted from eight states with levels surpassing EPA-designated MCLs. Four heavy metals (sodium, strontium, nickel, vanadium) were detected in samples submitted from ten states with levels surpassing EPA health guidance levels. Given that MCL and guidance levels represent limits of contamination protective of health, associated risks of consumption likely apply to both dogs and their owners. In this paper, we have made a point to include relative human demographic data alongside dog demographics (Table 1) in an effort to promote this practice as standard for future studies that aim to understand this sentinel relationship between dogs and their owners.

In addition to metals being identified at potentially unsafe levels, several of the responses to the water source survey warrant further investigation. For example, there was a positive correlation between water reported to be “soft” and the presence of iron identified in this study. Typically, if water softeners are used to ameliorate the build-up of minerals from heavy metals that cause water to be “hard” (e.g., magnesium and calcium), iron is also removed [47]. Furthermore, the differences in metals correlated with reportedly “soft” water, and those with the reported use of water softener is odd. Some of these findings may be complicated by even higher pre-treatment levels of a metal such as iron. On the other hand, the higher average values of several metals in water samples from older wells compared to newer wells (which correlated negatively with values of silicon, zinc, and lead) is reasonable given technological advances regarding pipe and tank materials.

Due to the multifactorial nature of many of the reported health conditions and the cross-sectional nature of this evaluation, causal relationships between water quality and health outcomes cannot be established from these data, despite instances of statistical significance. Specifically for metals already known to be dangerous at certain levels (e.g., titanium and chromium), some of the observed statistical associations are quite implausible in real-world scenarios. Nevertheless, observed associations between dogs’ health outcomes and the presence of heavy metal concentrations above EPA-recommended limits in more than one drinking water sample warrant further investigation. Additionally, the stastically negative associations between reverse osmosis water systems (installed as an intended health intervention) and developed health conditions, compared to positive associations between installed sediment filters (an aesthetic only intervention) and developed health outcomes warrants further investigation of the real-world efficacy of point-of-use water treatment options. Analyses with larger groups of dogs will provide greater statistical power and confidence, enabling more accurate assessments of specific metal levels, treatments, and specific outcomes such as the presence or absence of various health conditions (e.g. cancers, cardiac issues, gastrointestinal disease, etc.). With prospective studies rather than this cross-sectional design, causal analyses can be conducted.

For example, the addition of sodium compounds is regularly used during water treatment and softening and can increase sodium levels to more than 300 mg/L in drinking water, per the EPA [48]. However, even with increased sodium levels (which could adversely impact health [49]), treated water may contribute to better overall health outcomes in dogs compared to those drinking untreated groundwater. It could also be the case that individual owners who are concerned about implementing proper water treatment are likewise prone to maintaining their dog’s health through regular veterinary care.

Continued examination of the presence of several of the metals found in dogs’ drinking water is of utmost importance to evaluating and addressing water-related toxicant exposures. The known toxicity of environmental lead exposure, alone, continues to be a serious public health concern. Chen et al. (2023) [37] report that the blood lead levels of certain mammals, particularly dogs, are correlated with the levels of humans living in the same environment.

Furthermore, regionality may be especially important to identifying and understanding sources of heavy metal poisoning. Forte et al. (2023) [50] found detectable concentrations of both essential and nonessential metal(loids), including known endocrine-disrupting toxicants cadmium and lead, in the ovarian tissues of free-ranging female dogs in Italy. Toyomaki et al. (2020) [51] analyzed blood lead (Pb) concentrations and biochemistry of 120 domestically owned dogs living around a lead mining area in Kabwe, Zambia, and found that Pb isotope ratios in dogs decreased with increasing age and distance from the mine, and were similar to those previously reported in humans at the location.

While lead is a clearly identified culprit in severe adverse health effects, less is understood about the potential impacts of many other metals and elements, including those we have identified here in dogs’ drinking water. The additional and potentially amplifying impacts of contamination for certain communities and individuals (e.g., those with fewer resources, limited access to safety measures) is especially concerning. On the other hand, the presence of certain elements in drinking water may have ameliorating effects and should be incorporated into treatment. Given that our total number of sample returns with metals above EPA limits were relatively low in this sample group (excepting sodium) [30,42–46], a larger sample will be necessary to detect statistically significant differences among participants. Thus, we will deploy a second batch of collection kits to a larger, stratified sample of dogs (rural vs. urban X large vs. small), which will also offer the potential for case-control study design approaches. We will compare these data with previously collected data on dogs’ known health conditions to determine if, when, and where water quality correlates with health outcomes, with attention to age-specific risk.

## Conclusions

Drinking water toxicity from heavy metals and other contaminants can lead to both acute and chronic health conditions including organ failure in dogs [15,16]. Pet dogs, who have little awareness of or control over the quality of their drinking water, may be especially susceptible to such risks [32,33]. Furthermore, because previous and emerging research supports dogs’ role as sentinels of human health and wellbeing [36,52,53], heavy metal toxicity of drinking water should be regarded as a concern for humans living in the same households and homesteads.

## Institutional Review Board

The University of Washington IRB deemed this work to be human subjects research that qualifies for Category 2 exempt status (IRB ID no. 5988, effective 10/30/2018). All study-related procedures involving privately owned dogs were approved by the Texas A&M University IACUC, under AUP 2021-0316 CAM (effective 12/14/2021).

## Acknowledgments

We would like to acknowledge and thank the DAP participants who took the time to participate in our study and Jeffrey Parks for his assistance in completing the laboratory portion of this study.

## Dog Aging Project Consortium authors

Joshua M. Akey, Brooke Benton, Elhanan Borenstein, Marta G. Castelhano, Amanda E. Coleman, Kate E. Creevy, Kyle Crowder, Matthew D. Dunbar, Virginia R. Fajt, Annette L. Fitzpatrick, Unity Jeffery, Erica C Jonlin, Matt Kaeberlein, Elinor K. Karlsson, Kathleen F. Kerr, Jonathan M. Levine, Jing Ma, Robyn L McClelland, Daniel E.L. Promislow, Audrey Ruple, Stephen M. Schwartz, Sandi Shrager, Noah Snyder-Mackler, M. Katherine Tolbert, Silvan R. Urfer, Benjamin S. Wilfond

## Notes

https://dogagingproject.org/data-access

### Competing Interest Statement

The authors have declared no competing interest.

