## Supplementary Materials for "Testing for heavy metals in drinking water collected from Dog Aging Project participants"

---

Survey Title: Water Quality

Page header: About the Water Quality Survey

This survey is designed to collect more detailed information about your dog's primary drinking water source. You will have the opportunity to describe the water source from which your dog drinks and tell us about other water use in relation to your dog as well.

There may be some repetition between this survey and previous Dog Aging Project surveys; this is intentional. Overlap is part of our research design and is intended to aid us in corroborating information from different data sources.

The purpose of this survey is to gain an enhanced understanding of the quality of the water your dog is drinking. This is a very important aspect of canine health, and we want to be sure we understand the mechanisms of how it impacts health. Your help in completing this survey is greatly appreciated and will help us continue to learn more about the aging process in our canine companions.

Next Page >>

---

Page header: Your Primary Water Supply

**What is the primary water supply source for your household?**

1, Municipal
2, Non-municipal well water (treated/filtered)
3, Non-municipal well water (untreated/unfiltered)
4, Bottled or commercially delivered
5, Spring water
6, Rainwater cistern
98, Other
99, Don't know

If Other, ask:

**Please describe the primary water supply source for your household:** [open text]

If well water (2 or 3), ask:

**Which of the following best describes your well**:

1, Drilled well
2, Dug or bored well
99, Don't know

**Do you know how deep your well is?**

1, Yes
0, No

If Yes, ask:

**How deep is your well measured in feet?** [whole number only]

**Do you know what year your well was constructed?**

1, Yes
0, No

If Yes, ask:

**Please enter the year your well was constructed:** [integer]

<< Previous Page Next Page >>

Save & Return Later

---

Page header: Your Household Water

**Is your primary water supply located within 100 feet of the following features? Please select all that apply.**

0, None
1, Septic drain field
2, Pit privy or outhouse
3, Home heating oil storage tank
4, Pond, river, or freshwater stream
5, Cemetery
6, Tidal shoreline, estuary, or marsh
99, Don't know

**Is your primary water supply located within a ½ mile of any of the following sites? Please select all that apply.**

0, None
1, Landfill
2, Golf course
3, Abandoned quarry or industrial site
4, Active quarry
5, Active industrial, manufacturing, or processing site
6, Dump site
7, Field crops or plant nursery
8, Farm animal operation
9, Railroad tracks
10, Airport
11, Fracking site
12, Commercial underground storage tank or supply lines (e.g., gas station)
99, Don't know

If 5, ask:

**Please specify the type of industrial, manufacturing, or processing site**: [open text]

**Is your yard treated to control weeds?**

5, Annually, with regularity
4, Seasonally, with regularity
3, Annually, sporadically

2, Seasonally, sporadically
1, Infrequently
0, Never

**Is your yard treated to control insects and pests?**

5, Annually, with regularity
4, Seasonally, with regularity
3, Annually, sporadically
2, Seasonally, sporadically
1, Infrequently
0, Never

**Are any of the following water treatment systems currently installed and functioning properly? Please select all that apply.**

0, None
1, Ultraviolet (UV) Light
2, Water Softener (Conditioner)
3, Sediment Filter
4, Reverse Osmosis
5, Iron Removal
6, Activated Carbon (Charcoal) Filter
7, Chlorination System
8, Acid Neutralizer
98, Other
99, Don’t know

If Other, ask:
Please specify the other type of water treatment system: [open text]

**Have you ever had your primary water supply tested?**

1, Yes
0, No

**What types of pipes do you have in your home?**

3, Copper
4, Lead

5, Galvanized steel
2, Plastic (PVC, PEX, etc.)
98, Other
99, Don’t Know

If Other, ask:
**Please specify other type of pipe material in your home**: [open text]

**Do you have problems with corrosion, pitting, or pinhole leaks in pipes or plumbing fixtures?**

1, Yes
0, No

**Do you consider your household water “hard” or “soft”?**

1, Hard
2, Soft
3, Neither
99, Don’t know

**Does your water stain plumbing, cooking appliances, utensils, or laundry?**

1, Yes
0, No

If yes, ask:

**How would you describe the stains? Please select all that apply.**

1, Blue or green
2, Rusty, orange, or brown
3, Black or gray
4, White or chalky
98, Other

If Other, ask:

**Please specify the color of the water stains**: [open text]

**Does your water have an unpleasant taste?**

1, Yes
0, No

If Yes, ask:

**How would you describe the taste of your water?**

1, Bitter
2, Rotten egg (sulfur)
3, Salty
4, Metallic
5, Oily
6, Soapy
98, Other

If Other, ask:

**What does your water taste like?** [open text]

**Does your water have an unpleasant odor?**

1, Yes
0, No

If Yes, ask:

**How would you describe the odor of your water?**

1, Rotten egg (sulfur)
2, Kerosene or gas
3, Musty
4, Chemical
98, Other

If Other, ask:

**What does your water smell like?** [open text]

**Does your water have an unnatural color or appearance?**

1, Yes
0, No

If Yes, ask:

**How would you describe the color or appearance of your water?**

1, Muddy
2, Milky
3, Black or gray tint
4, Yellow tint
5, Oily film
98, Other

If Other, ask:

**What does your water look like?** [open text]

<< Previous Page     Next Page >>

Save & Return Later

---

Page header: About Your Dog

**Would you say in general your dog's health is:**

1, Excellent
2, Very good
3, Good
4, Fair
5, Poor
6, Very poor

**Does your dog have any of the following ongoing health conditions? Please select all that apply.**

1, Infectious or parasitic disease
2, Ingestion of toxic or controlled substance
3, Trauma
4, Cancer or tumors
5, Eye disorders
6, Ear, nose, and throat disorders
7, Dental or oral disease
8, Skin disorders
9, Cardiac disorders
10, Respiratory disorders
11, Gastrointestinal disorders
12, Liver or pancreas disorders
13, Kidney or urinary disorders
14, Reproductive system disorders
15, Orthopedic disorders
16, Neurologic disorders
17, Endocrine disorders
18, Hematopoietic (blood or lymphatic) disease
19, Immune-mediated disease
98, Other disease or disorder
99, None

If Other, ask:

What other ongoing health conditions does your dog have? [open text]

**Are there other animals present in your home or on your property?**

1, Yes
0, No

If Yes, ask:

**What types of animals?**

Dogs
Cats
Birds
Reptiles
Livestock

Horses
Rodents
Fish
Wildlife
Other

If Other, ask:

**What other types of animals?** [open text]

**Does your dog drink out of any of the following water sources? Please select all that apply.**

1, Municipal
2, Non-municipal well water (treated/filtered)
3, Non-municipal well water (untreated/unfiltered)
4, Bottled or commercially delivered
5, Spring water
6, Rainwater cistern
7, Outdoor water source (e.g., pond, stream, puddles)
8, Toilet water
98, Other
99, Don't know

If Other, ask:
**Please describe the other water supply your dog drinks out of**: [open text]

**In your dog's water bowl, do you notice floating or settled particles?**

1, Yes
0, No

If Yes, ask:

**How would you describe these particles? Please select all that apply**.

1, White flakes
2, Black specks
3, Red-orange slime

4, Brown sediment
98, Other

If Other, ask:

What do the particles look like? [open text]

**In what ways do you use your household water for your dog? Please select all that apply.**

1, Drinking water
2, Brushing dog's teeth
3, Bathing dog
4, Washing bedding or toys
5, Washing food and water dishes
98, Other
99, None of these

If Other, ask:

**In what other ways do you use your household water for your dog?** [open text]

**Is there anything else that you think we should know about the various sources of water that your dog comes into contact with?** [open text]

**Would you be willing to collect a water sample from your dog's primary drinking water source for our water quality study using a free kit provided by the Dog Aging Project?**

1, Yes
0, No

---

If No to last question, use this Woof text as final page::

**Woof! Thank you for completing the Water Quality Survey!**

**Our research team will use the valuable information you've provided to enhance our understanding of the environmental exposures that your dog might experience.**

**Ultimately we want to know more about how water quality can affect our health. As our study progresses, we will keep you informed of what we learn.**

***Please click Submit below to finalize your answers and close this task*.**

<< Previous Page     Submit

---

If Yes to last question, use this Woof text as final page::

**Woof! Thank you for completing the Water Quality Survey and expressing interest in the next phase of this study!**

**In the next few weeks, we will be analyzing the information collected from this survey and identifying geographic regions from which we'd like to collect water samples. If you are in one of these regions, we will send you an email with more details, and you can accept or decline at that time.**

**If you choose to participate, we will send you a kit that contains a water collection container, printed instructions about how to collect the sample, and a prepaid mailer for returning the water sample to our lab. There will be no cost to you.**

**This is the mailing address that we have on file for you:**

[pipe in shipping address]

**Is this the best address for you to receive the Water Sampling Kit?**

1, Yes
0, No

If No, allow address correction:

Please note that changing your mailing address here will <u>not</u> change your primary address in your research portal.

**Thank you again for your participation! Our research team will use the valuable information you've provided to enhance our understanding of the environmental exposures that your dog might experience. Ultimately we want to know more about how water quality can affect our health. As our study progresses, we will keep you informed of what we learn.**

***Please click Submit below to finalize your answers and close this task.***

<< Previous Page Submit

**DOG AGING PROJECT WATER SAMPLING KIT**
***Read these instructions the day before filling your bottle!***

- Step 0. Plan ahead. It is ***very*** important to collect water **after allowing it to sit in your pipe for at least 6 hours**. To do so easily, water should be collected first thing in the morning (after no one uses the water from that tap overnight) *from your dog's primary drinking water source*.

- Step 1. After the water was not used for at least 6 hours, open the kit and remove the plastic cap from the bottle.

- Step 2. With the 250 mL collection bottle held under the water tap, open the cold water tap and fill the bottle at high flow (as if you were filling a pitcher of water).

- Step 3. Turn off the flow, put the cap back on the bottle, and firmly tighten the cap.

- Step 4. Complete the sample label as instructed above.

- Step 5. Place the bottle into the Ziplock bag that is provided. Seal the Ziplock bag and place the bag (with bottle inside) into the box that is provided.

- Step 6. Tape the box shut, apply the pre-paid return shipping label, and drop at any UPS collection site.

- Call the Ruple Lab at 540-231-0342 or if you have any questions regarding these instructions.

Dear {Participant},

Thank you for participating in our water quality study for the Dog Aging Project! We are so grateful for your help with this research and wanted to provide you with the results of our analysis.

After we received your sample, we analyzed [Dog's name]'s drinking water for the presence of 28 different elements. Many of these elements are commonly found in drinking water and are not known to cause any health issues. However, eight of the elements are heavy metals that are listed by the Environmental Protection Agency (EPA) as having Maximum Contaminant Level Goals (MCLGs).

If one of these metals occurs in your water at a level below the MCLGs, the EPA considers the water safe to drink. The EPA is concerned about levels above the MCLGs and provides additional information and precautionary steps to consumers like yourself at this link: National Primary Drinking Water Regulations | US EPA.

Below, we display the results for this study for the eight heavy metals in parts per billion (ppb). In the table, we present two numbers for each heavy metal:

- *MCLG: EPA Maximum Contaminant Level Goal for each metal.*
- *YOUR RESULT: the level from the water sample you submitted.*

The MCLGs for each metal are depicted as the black vertical line. Your result is the small blue line.

| | | MCLG | **YOUR RESULT** |
|---|---|---|---|
| Arsenic | 0 – 20 ppb | 10 | 0 |
| Barium | 0 – 4000 ppb | 2000 | 2.5 |
| Cadmium | 0 – 10 ppb | 5 | 0 |
| Chromium | 0 – 200 ppb | 100 | 0 |
| Copper | 0 – 2600 ppb | 1300 | 1415.4* |
| Lead | 0 – 30 ppb | 15 | 0.9 |
| Selenium | 0 – 100 ppb | 50 | 0 |
| Uranium | 0 – 60 ppb | 30 | 0 |

MCLG **YOUR RESULT**

At least one of your values (denoted in red with an asterisk) is to the right of the MCLG line. We recommend that you visit National Primary Drinking Water Regulations | US EPA for more information.

We also tested 20 other elements that are not regulated by the EPA. Not much is known about the effects of these other 20 elements in ______'s drinking water. This current lack of information is part of why we are studying this (with your help)! The table below lists each element and displays the following results in either parts per million (ppm) or parts per billion (ppb). In the absence of known MCLGs for these elements, we have also provided the quartile your result is in compared to other study participants to allow for some comparison.

| | Quartile | | | | **YOUR RESULT** | | Quartile | | | | **YOUR RESULT** |
|---|---|---|---|---|---|---|---|---|---|---|---|
| | 1 | 2 | 3 | 4 | | | 1 | 2 | 3 | 4 | |
| Aluminum | * | | | | 0.0 ppb | Potassium | * | | | | 0.0 ppb |
| Calcium | * | | | | 240.0 ppb | Silicon | | | * | | 11704.1 ppb |
| Chlorine | * | | | | 3.1 ppm | Silver | * | | | | 0.0 ppb |
| Cobalt | * | | | | 0.0 ppb | Sodium | | | * | | 38687.6 ppb |
| Iron | * | | | | 0.0 ppb | Strontium | * | | | | 0.0 ppb |
| Lithium | | | | * | 11.2 ppb | Sulfur | | * | | | 10.6 ppm |
| Magnesium | | * | | | 228.0 ppb | Tin | * | | | | 0.0 ppb |
| Manganese | | | | * | 5.5 ppb | Titanium | | | | * | 1.3 ppb |
| Nickel | | | | * | 5.8 ppb | Vanadium | * | | | | 0.0 ppb |
| Phosphorus | | * | | | 5.4 ppb | Zinc | | | | * | 470.5 ppb |

Quartile 1: Your result is in the lowest 25% of study participants.
Quartile 2: Your result is higher than 25% and lower than 50% of study participants.
Quartile 3: Your result is higher than 50% and lower than 25% of study participants.
Quartile 4: Your result is in the highest 75% of study participants.

If you have any additional questions about your results, consider reaching out to your local community health department. If you have questions about this study, please. Thank you again for your participation in this important study!

Sincerely,

The team at the Dog Aging Project

Dog Aging Project

Longer, healthier lives. Together.
